# Hydrostatic pressure and lateral actomyosin tension control stretch and tension of the basement membrane in epithelia

**DOI:** 10.1101/2022.09.02.506324

**Authors:** Karla Y. Guerra Santillán, Christian Dahmann, Elisabeth Fischer-Friedrich

## Abstract

The shaping of epithelial tissues into functional organs often depend on asymmetries in mechanical tension present at the apical and basal sides of cells. Contraction of an actomyosin meshwork underlying the apical side of cells is known to generate apical tension. The basal side of cells is also associated with an actomyosin meshwork, but it is, in addition, connected to a specialized extracellular matrix, the basement membrane. However, how basal tension is generated, and the role of the basement membrane in this process, are not well understood. Here, using atomic force microscopy, we measure mechanical tension in the basal surface of the wing disc epithelium of *Drosophila*. We find that basal tension depends on both the actomyosin cytoskeleton and the basement membrane, and that it is proportional to lateral surface tension and hydrostatic pressure. Collagen IV turnover and mobility are slow indicating that the basement membrane can store elastic stresses. Our data suggest that elastic stresses in the basement membrane induced by basement membrane stretch are a key factor in the adjustment of basal tension. Hydrostatic pressure and lateral actomyosin contractility are two driving forces by which epithelial cells can maintain this basement membrane stretch.

## I. INTRODUCTION

In animal development, morphogenesis is the process by which cells and tissues undergo a series of deformations and translocations to attain their final form. During morphogenesis, some cells organize in sheet-like structures known as epithelial tissues. Epithelial morphogenesis typically includes a series of epithelial folding events that generate a complex three dimensional structure. This process depends on the generation of mechanical force and tissue compliance^1^. Failure to properly reshape epithelia during development can lead to severe birth defects providing evidence that emergence of proper organ form is an important prerequisite for organ function^2,3^.

Epithelia consist of cells which are polarized along their apical-basal axis. Neighboring cells are mechanically connected at their sub-apical regions through adherens junctions that are linked to the actomyosin cytoskeleton. The basal surface of cells is connected to a basement membrane composed by extracellular matrix (ECM) proteins^4^. The composition of basement membranes is to a great degree preserved across the animal kingdom^5^ and includes Laminin, Collagen IV, Nidogen, Perlecan and Agrin^6^. Out of these, Collagen IV was shown to be particularly crucial for the mechanical integrity of the basement membrane^7–9^. The basement membrane is important in the morphogenesis of epithelia, for example in the branching of the lung epithelium and the mammary gland epithelium^10,11^ as well as in egg chamber elongation in *Drosophila*^7,12^. Overall, epithelial morphogenesis relies on asymmetries in epithelial mechanics such as differences in apical and basal tension^13^. Therefore, it is of vital importance to understand the regulation of both apical and basal tension. Research on the mechanical forces that drive epithelial sheet movements has been mainly focused on the generation of apical tension by the actomyosin cytoskeleton^14^. By contrast, the specific contributions of the basement membranes in the generation of basal tension in epithelial tissues remain unclear^15^.

The developing *Drosophila* wing is a useful model to understand how the mechanical properties of the basement membrane contribute to epithelial morphogenesis. The wing develops from a larval precursor epithelium, the wing imaginal disc. The wing imaginal disc is a single-cell layered epithelium forming a flat sac-like structure with the apical surfaces facing the interior. Cells on the one side of wing imaginal disc are squamous-shaped, whereas cells on the opposing site are columnar. The columnar epithelium gives rise to the future wing blade, notum and hinge tissue of the adult fly^16^. The basement membrane of the wing imaginal disc plays important roles during the morphogenesis of this tissue. For example, the highly columnar shape of cells and the overall bending of the wing imaginal discs towards its basal side depend both on an intact basement membrane^17–19^. Moreover, the formation of a fold in the center of the prospective hinge region involves the local reduction of ECM density and a decreased basal tension^19^. However, how basal tension is generated in wing imaginal discs remains to be elucidated.

Here, we quantify basal tension in the wing disc epithelium using atomic force microscopy and present a quantitative analysis of how basal tension is generated. We find that the actomyosin contractility and the basement membrane are required for the generation of basal tension. Using live-imaging of explanted wing discs, we show that Collagen IV turnover and mobility in the wing disc is slow indicating solid-like material properties on the time scale of epithelial folding. Perturbing lateral actomyosin contractility and intracellular hydrostatic pressure, we change the epithelial basal area and find evidence that elastic stretch in the basement membrane makes a considerable contribution to basal tension in the wing disc epithelium. This work reveals further how active molecular force generation in epithelial cells is relayed to passive mechanical stresses in the basement membrane.

## II. RESULTS

### A. Basal tension depends on the basement membrane and on the actomyosin cytoskeleton

To quantify basal tension, we performed shallow indentations into the basal surface of explanted wing discs with the pyramidal indenter of the cantilever of an atomic force microscope (AFM), see Fig. 1A,B and Materials and Methods. As a readout, we obtained force-indentation curves, see Fig. 1C, showing the rise of measured AFM force *F* in dependence of cantilever tip indentation *δ*. Notably, force-indentation curves were observed to be approximately linear, see Fig. 1C and Fig. S2. Applying a previously established analysis scheme for the measurement of interfacial cell tension^20,21^, we estimated basal tension in the wing disc as 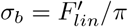 throughout this manuscript, where 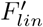 is the fitted linear slope of the measured force-indentation curve, see Fig. 1C and Materials and Methods. Interestingly, obtained basal tension values were in the range of 0.3 – 0.5 mN/m for wing discs explanted 72 h after egg lay (AEL), see Fig. 1D-G. Therefore, tension values closely matched values of actin cortical tension measured in interphase cells via micropipette aspiration or AFM measurements^22–25^.

**FIG. 1.**
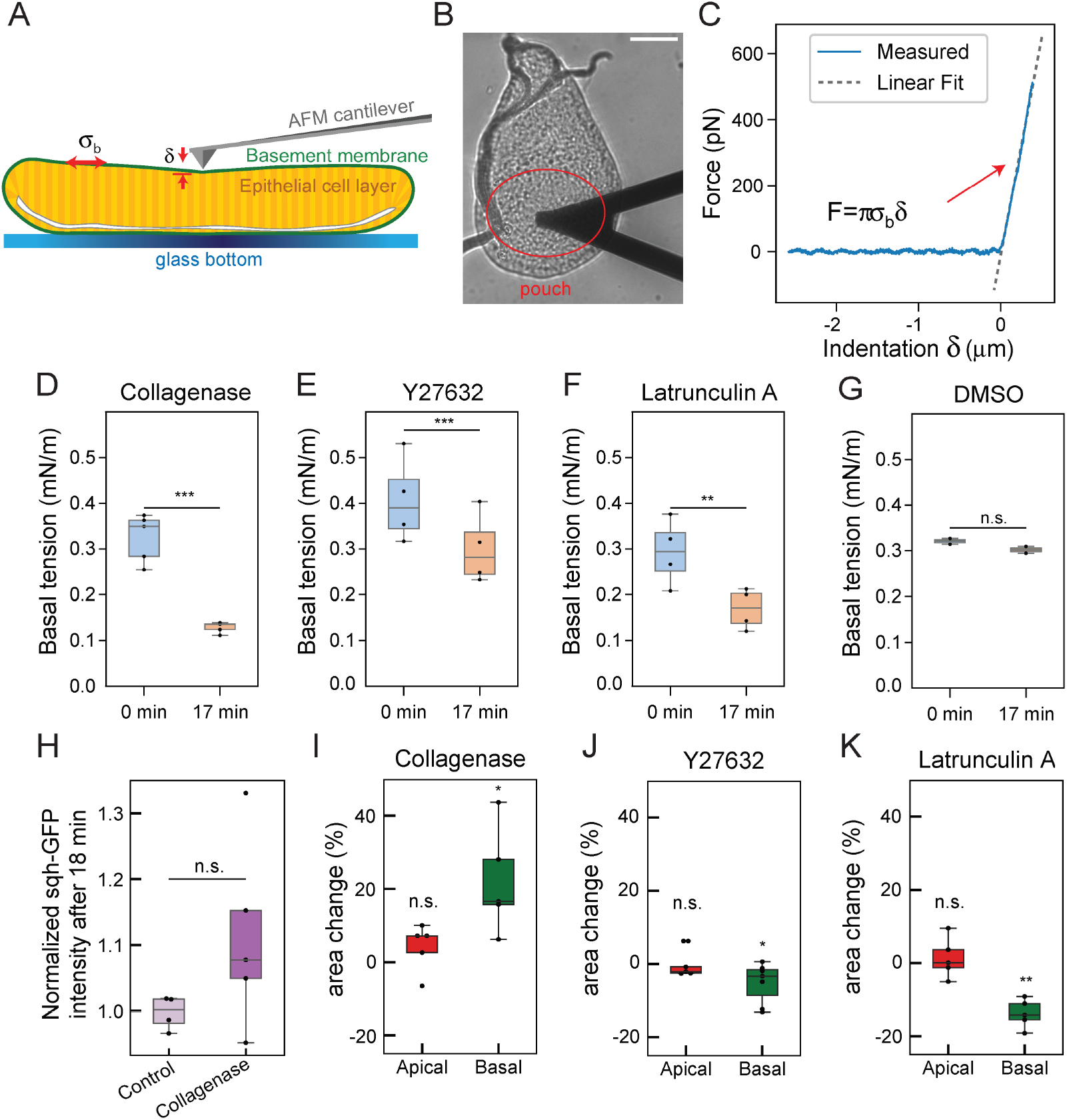
Basal tension depends on the basement membrane and on the actomyosin cytoskeleton. A) Schematic illustrating the measurement of basal tension (*σ_b_*) of wing discs. The force applied by an AFM cantilever results in the indentation (*δ*) of the basal surface. B) Brightfield micrograph of an explanted wing disc during an AFM measurement. Scale bar: 50 *μ*m. C) Exemplary force-indentation curve recorded by the AFM during the lowering of the cantilever over the tissue. The force *F* starts to increase approximately linearly after the cantilever has touched the tissue surface. A linear fit is shown by the dashed line. The magnitude of the linear slope divided by π is used as an estimate of basal tension. D-G) Boxplots showing basal tension estimates measured in the central pouch region of 72 h AEL wing discs before (0 min) and after 17 min incubation with indicated reagents (Collagenase: 0.2 mg/ml, Y27632: 1 mM, Latrunculin A: 4 μM, DMSO: 2%, N=4 wing discs for each treatment, DMSO: N=2). H) Boxplots showing the changes of Myosin II in the pouch at the basal side judged by normalized Sqh-GFP fluorescence intensity after 18 min of incubation in control conditions and with Collagenase (0.2 mg/ml). The fluorescence intensity was normalized by the initial value of each wing disc (Control: N=4, Collagenase: N=5, same concentrations as in D and I). I-K) Boxplots indicating changes of apical and basal cell areas upon 17 min incubation with indicated reagents (Collagenase, Latrunculin: N=5, Y27632: N=7, same concentrations as D-F). Significance was determined with a paired T-test (D-G,I-K) or a Mann-Whitney U-Test (H) (*p < 0.05, **p < 0.01, ***p < 0.001, n.s. p> 0.05 not significant).

Basal tension may depend on the basement membrane and/or the actomyosin cytoskeleton. To test this hypothesis, we first explanted wing discs 72 h AEL and treated them with Collagenase, which degrades basement membrane components^7^. Collagenase treatment resulted in a reduction of basal tension by ≈ 70% (Fig. 1D and Supplementary Fig. S1A). Next, we perturbed the actomyosin cytoskeleton. Treatment of explanted wing discs with the Rock inhibitor Y27632 reduced basal tension by ≈ 30 % (Fig. 1E and Supplementary Fig. S1B). Co-incubation with Latrunculin A - an inhibitor of actin polymerisation - resulted in a reduction of basal tension by 50 % (Fig. 1F and Supplementary Fig. S1C)). As a control experiment, we verified that incubation of wing discs in 2% DMSO did not significantly change basal tension (Fig. 1G).

In order to test whether tension reduction upon Collagenase treatment was due to a decrease in actomyosin contractility mediated by ECM degradation, we monitored the time evolution of fluorescently labeled Myosin II (Sqh-GFP) at the basal side upon addition of Collagenase. After 18 min of co-incubation, we find on average a slight increase of Myosin II at the basal side indicating that actomyosin contractility is slightly elevated, see Fig. 1H and Supplementary Fig. S1D-F. This data suggest that the observed reduction in basal tension upon Collagenase treatment is not caused but, if anything, slightly mitigated by accompanying changes in actomyosin. Taken together, we conclude that the actomyosin cytoskeleton and the basement membrane contribute both to the generation of basal tension.

In addition, we monitored the change of apical and basal cell areas in response to treatments with the above-mentioned drugs. None of the reagents changed apical cell areas significantly. For basal cells areas, we interestingly found opposite changes in response to Collagenase and actomyosin perturbing drugs. While Collagenase treatment led to a median increase of basal cell areas of ≈ 16%, the Rock inhibitor Y27632 and Latrunculin A both reduced basal cell areas by 3% and 13%, respectively, see Fig. 1I-K and Supplementary Fig. S1G-I. We conclude that tension reduction upon Collagenase-induced basement membrane degradation is linked to basal widening. On the other hand, basal tension reduction through deactivation of actomyosin contractility is connected to basal shrinkage.

### B. Basal tension depends on developmental time and wing disc position

The spatiotemporal control of mechanical tension contributes to epithelial morphogenesis^26–28^. We therefore next tested whether basal tension depended on developmental time and/or position within the wing disc. Wing discs are relatively flat structures until a developmental time point of ≈ 72 h AEL. Thereafter, three stereotypic folds form in the hinge region of the wing disc, see Fig. 2A. We therefore probed basal tension in a developmental time window around fold initiation at 64, 68, 72, 76, and 80 h AEL (H/H fold formation begins approximately at 72 h AEL). For this analysis, we measured basal tension in the pouch (which gives rise to the adult wing blade), because, once the tissue was folded, tension measurements in the hinge region were not possible anymore. Basal tension was relatively low at 64 h AEL, significantly increased by 68 h AEL, and thereafter declined to reach a value at 76 and 80 h AEL similar to the one at 68 h AEL, see Fig. 2B. Thus, basal tension peaks right before the onset of H/H fold formation and thereafter relaxes to a value similar as before fold formation. Interestingly, previous work reported an increase of basal edge tension in the wing disc pouch at a similar time point in development, further illustrating the reconfiguration of internal stresses in this time window^19^.

**FIG. 2.**
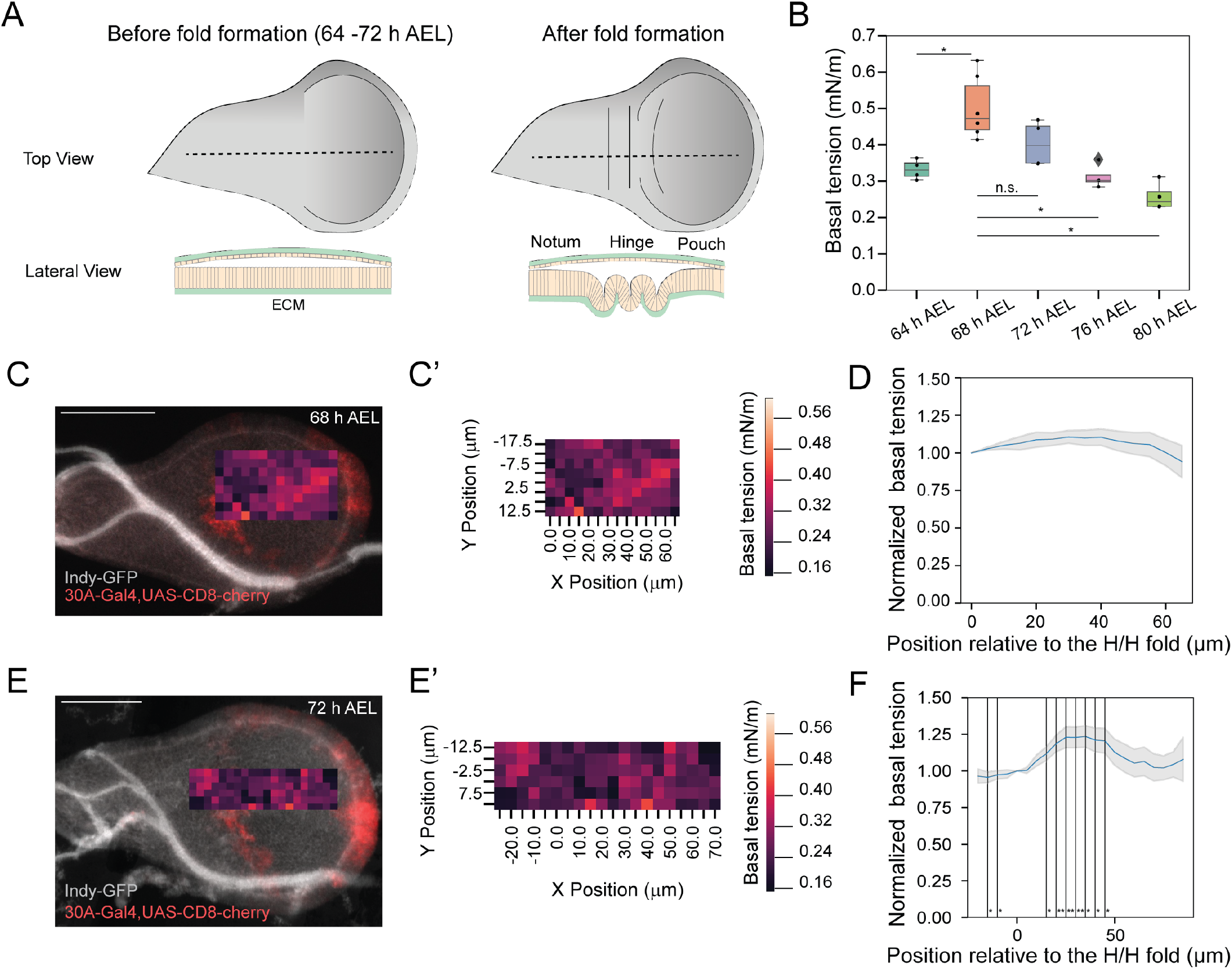
Basal tension depends on developmental time and wing disc position. A) Schematics of top views and lateral views of wing discs before and after fold formation. B) Boxplot of basal tension in the pouch center at the indicated stages of development. 64 h AEL (n=4 wing discs), 68 h AEL (n=5), 72 h AEL (n=4), 76 h AEL (n=6) and 80 h AEL (n=4). Significance was determined with a Mann-Whitney U-Test (*p< 0.05, **p< 0.01, ***p< 0.001, n.s. not significant). C-F) Basal tension measured across different regions of the wing disc at 68 h AEL (C-D) and 72 h AEL (E-F). C,E) Confocal image of measured wing discs expressing Indy-GFP (gray, the bright structures are tracheae) and 30A-Gal4,UAS-CD8-cherry (red). The latter labels prospective H/H fold cells. Scale bars: 50 *μ*m. Heatmaps show the basal tension measured in a 65 *μ*m × 25 *μ*m (C) and a 100 *μ*m × 25 *μ*m (E) grid along the wing disc, respectively. C’, E’) Heat maps from (C) and (E) with quantitative scales. D, F) Mean and s.e.m. normalized basal tension along the wing disc long axis (X-axis) relative to the prospective H/H fold cells for n=7 and n=11 wing discs, respectively. Tension was normalized by the tension value of the prospective H/H fold cells (X = 0 position). Significance was determined with a Mann-Whitney U-Test (*p< 0.05, **p< 0.01, ***p< 0.001, n.s. not significant).

Next, we investigated the dependence of basal tension on the position within the wing disc. To this end, we performed a series of indentations along a spatial grid covering a region from the hinge to the pouch at two different developmental time points, before (68 h AEL) and at the onset of H/H fold formation (72 h AEL), see Fig. 2C,E. To align the measurements from different wing discs, the grids were positioned relative to the prospective H/H fold cells, as identified by the activity of the 30A-Gal4 line (RRID: BDSC_37534). At 68 h AEL, basal tension was similar for all positions analyzed. By contrast, at 72 h AEL, basal tension was significantly higher (by ≈ 25%) in the pouch region compared to the hinge region see Fig. 2D,F. Interestingly, a prior report showed that basal edge tension is higher in the pouch compared to the hinge region of the wing disc^19^. Taken together, our data demonstrate that basal tension depends both on developmental time and position within the wing disc.

### C. Long term imaging of Collagen IV suggests solid-like properties of the basement membrane

We next explored the physical mechanisms that contribute to basal tension in the wing disc. Based on our observation that basal tension depends critically on Collagen IV and, thus, on the integrity of the basement membrane, we hypothesized that basal tension could be, at least in part, generated through elastic stretch of the basement membrane. However, this hypothesis relies on a dominantly elastic, solid-like behavior of the basement membrane as otherwise stresses due to stretch would be quickly dissipated^29^.

To test this hypothesis, we explored the turnover of the basement membrane material, since viscous relaxation due to turnover of constituents is a common theme in the mechanical properties of living materials^24,30^. As Collagen IV is produced in the larval fat body^17^, i.e. outside of the wing disc, collagen turnover in explanted wing discs can be quantified by measuring Collagen IV abundance over time. We measured Collagen IV using the ColIVα2::GFP exon trap line (hereafter ColIV-GFP)^31^. Wing imaginal discs were explanted at 72 h AEL and ColIV-GFP was imaged for 7 hours at the basal side of the pouch and notum/hinge regions, see Fig. 3A-E. ColIV-GFP intensity decayed by ≈ 20 % in both pouch and notum/hinge region, see Fig. 3C,D. Fitting an exponential decay function to ColIV-GFP intensity over time, we found characteristic decay times above 10 hours, see Fig. 3D,E. To further test the mobility and turnover of Collagen IV in the wing disc basement membrane, we also performed fluoresence recovery after photobleaching (FRAP) experiments on the basement membrane of explanted wing discs expressing ColIV-GFP, see Fig. 3F-H. Longterm imaging of photobleached areas revealed that there was very little or no recovery of ColIV-GFP fluorescence (below 5% in seven hours after belaching, see Fig. 3H). In addition, the bleached fluorescence intensity profile stayed largely intact over seven hours indicating that there was little or no mobility of Collagen IV in the basement membrane (Fig. 3I). Taken together, our experiments of long-term imaging and photobleaching of ColIV-GFP in the basal side of the wing disc indicate that the turnover and mobility of Collagen IV in the basement membrane is slow as compared to the time scale of fold formation, i.e. few hours^19^. Therefore, we conclude that from the perspective of epithelial folding in the third instar larva, the basement membrane can be considered as a mostly solid-like material.

**FIG. 3.**
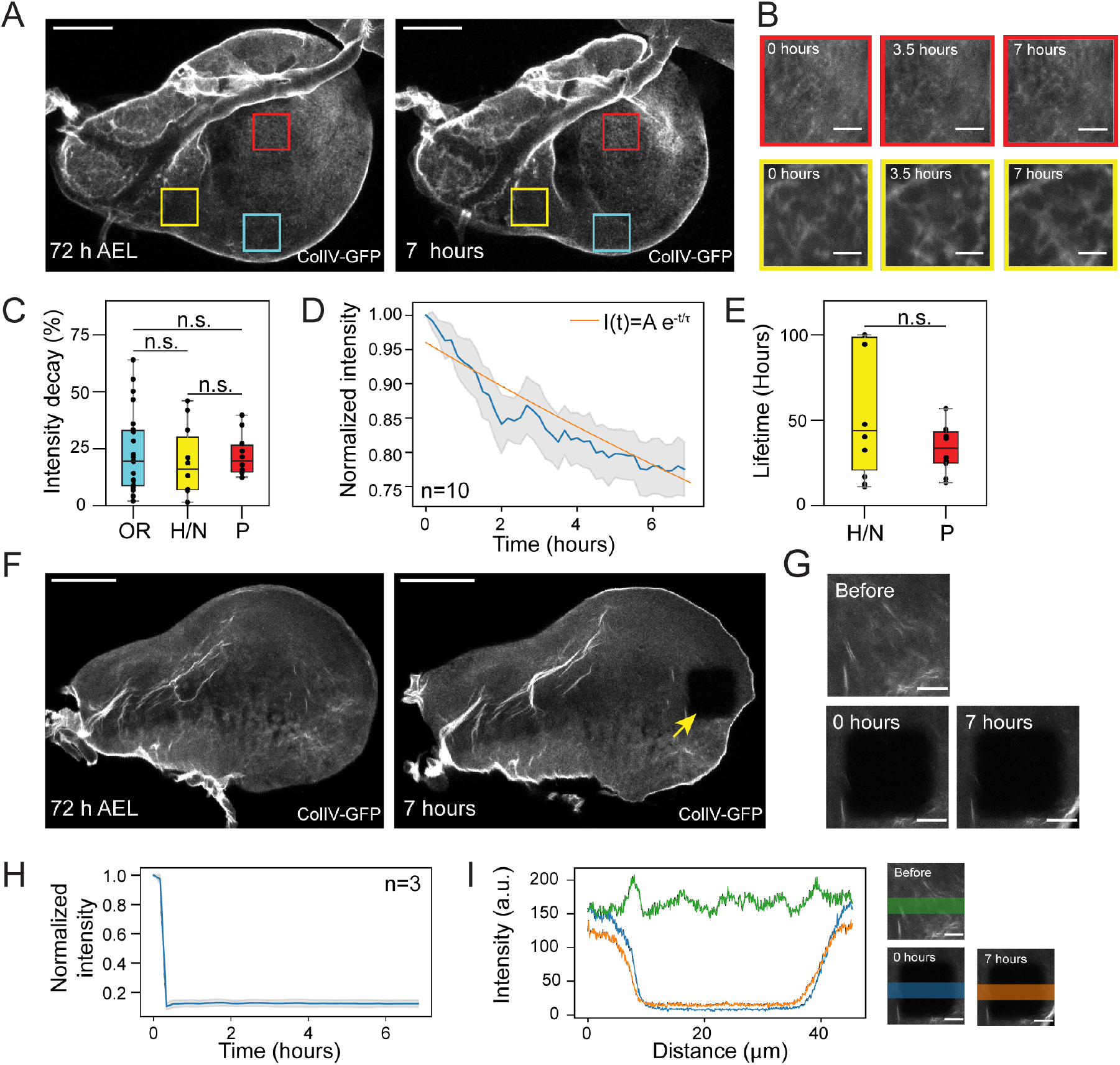
Long term imaging of Collagen IV suggests solid-like properties of the basement membrane. A) Sum fluorescence intensity projections of a Z-stack of images of the basal side of a 72 h AEL wing disc expressing ColIV-GFP freshly mounted (left panel) and 7 hours in culture. The regions indicated by the yellow and red 30 × 30 *μ*m squares were imaged every 10 min for 7 hours (yellow: hinge or notum (H/N), red: pouch (P)). The cyan 30 × 30 *μ*m square (OR) was only imaged at t=0 h and 7 h as a photobleaching control. Scale bars: 50 *μ*m. B) Magnified views of the regions indicated in (A) by the red and yellow squares at the indicated time points. Scale bars: 10 *μ*m. C) Boxplot of ColIV-GFP fluorescence intensity decay between t= 0 h and t= 7 h in different regions of the wing disc as indicated in panel A. D) Normalized ColIV-GFP fluorescence intensity over time of pouch-located 30 × 30 *μ*m squares averaged over ten wing discs (blue line). The fluorescence intensity was normalized by the initial value of each wing disc. The grey area indicates the standard error of the mean. An exponential decay is fitted (orange line) with decay time *τ* =27.2 *h*. E) Boxplot of lifetime of ColIV-GFP in the pouch (P) and hinge/notum (H/N) regions of wing discs (n=10 wing discs, each). F) Sum fluorescence intensity projections of a Z-stack of images of the basal part of a 72 h AEL wing disc expressing ColIV-GFP. Left panel: freshly mounted wing disc before photobleaching, right panel: 7 hours after photobleaching. The yellow arrow indicates the position of the 30 × 30 *μ*m square that was bleached and subsequently imaged every 10 minutes. Scale bars: 50 *μ*m. G) Images of the bleached area before bleaching and at time points t= 0 h and t=7 h, respectively. Scale bars: 10 *μ*m. H) Normalized mean ColIV-GFP fluorescence intensity over time after photobleaching averaged over three wing discs (blue line). The grey area indicates standard error of the mean. Normalized as in (D). I) ColIV-GFP fluorescence intensity along a vertical line scan overlapping a photobleached region (see panels on the right) before bleaching and at time points t= 0 h, and t=7 h, respectively. Significance was determined with a Mann-Whitney U-Test. n.s. not significant.

### D. A model of cellular force balances

Given our finding of a solid-like nature of the basement membrane, elastic stretch of the basement membrane should lead to an increase of basal tension by a passive stretch-induced contribution 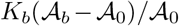 in addition to an active actomyosin-based contribution 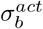. Here, 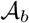 is the basal cell surface area and 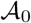 is the resting basal cell surface area in the absence of stress. *K_b_* is an elastic area bulk modulus of the basal cell surface.

Given this mechanical insight, we wanted to investigate the active processes and the mechanical driving forces in the cell that generate passive elastic stretch in the basement membrane. To this end, we considered an energy functional

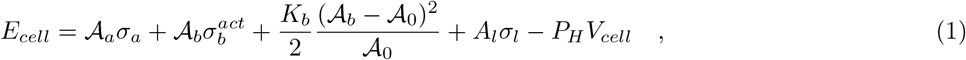

which is as a simplified version of the vertex model energy functional^32^ where *V_cell_* and *P_H_* denote cellular volume and cell-internal hydrostatic pressure *P_H_*, respectively. Here, we have included cell surface tensions for all cellular faces (apical: *σ_a_*, basal: *σ_b_*, lateral: *σ_l_*) with associated apical, basal and lateral surface areas 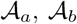 and *A_l_*. In the following, we assume a top-down-symmetric epithelium before fold formation with equal apical and basal cell surfaces 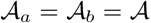, and identical apical and basal cell surface tensions *σ_a_* = *σ_b_*. By variation of the energy functional (1) with respect to 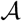 and *ℓ*, we obtain as conditions of minimal energy

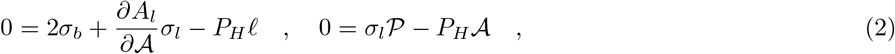

where we have used that 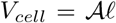, 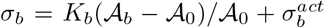 and that 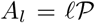 with 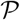 the perimeter of the apical and basal cell cross-section. For a hexagonal apical and basal cell cross-section with edge length *w*, we have 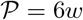 and 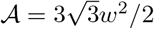 and thus 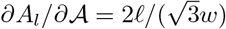. Consequently, we obtain from Eqn. (2)

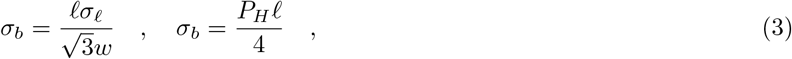

which corresponds to the force balance illustrated in Fig. 4. Therefore, basal and lateral tension are predicted to be proportional to each other, where basal tension is expected to be much larger than lateral tension due to the large aspect ratio *ℓ/w* of epithelial cells. Furthermore, basal and lateral tension are predicted to be proportional to hydrostatic pressure in the cell.

**FIG. 4.**
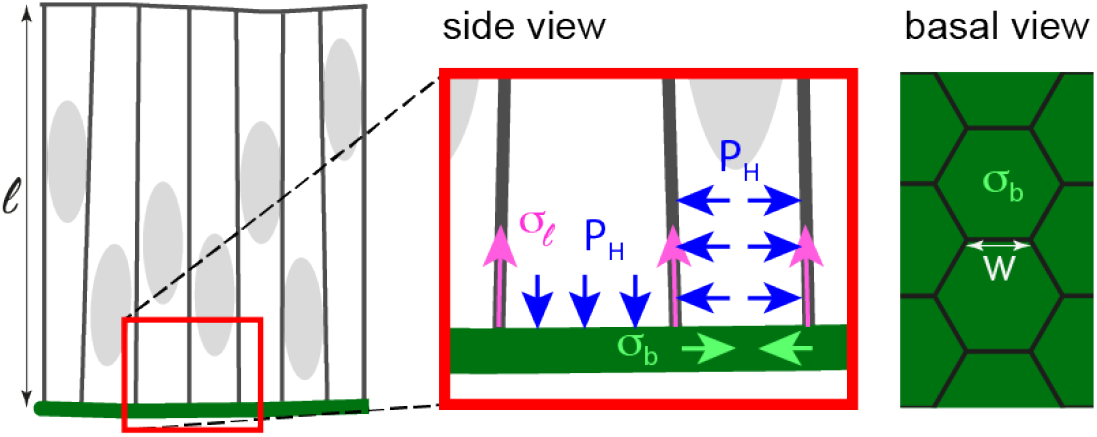
A model for vertical and horizontal force balance of the wing disc epithelium taking into account lateral, apical and basal tension as well as intracellular hydrostatic pressure. At the basal side, vertical force balance is achieved through i) hydrostatic pressure *P_H_* pushing against the basement membrane (blue arrows shown in left cell) and ii) the vertical component of contractile tension *σ_ℓ_* in the lateral cell surfaces pulling on the basement membrane (pink arrows on lateral cell interfaces). Horizontal force balance emerges through the interplay between i) contractile apical and basal tension (*σ_b_*, light green arrows) and ii) hydrostatic pressure pushing against the lateral cell surfaces in the epithelium (blue arrows shown in right cell). While lateral and apical tension are dominated by actomyosin activity, basal tension might stem from a combination of actomyosin contractility and elastic stretch in the basement membrane. Vertical and horizontal force balance require that 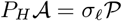 and that 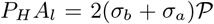, where 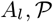 and 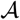 denote lateral cell surface area, the perimeter of the cellular cross section and the apical/basal cell surface area, respectively, see Eqn. (2) and (3).

In a top-down symmetric epithelium with hexagonal cross-section, for small changes of hydrostatic pressure *δP_H_* or lateral tension *δσ_ℓ_*, induced changes of basal tension are given to linear order by

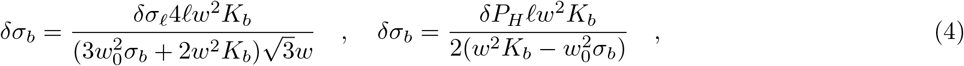

where we assumed that *δσ_a_* = 0 because *σ_a_* is subject to active regulation. In addition, we assumed constant cell volume such that 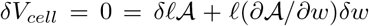 and therefore 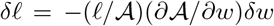. Furthermore, we anticipated that basal tension changes due to elastic stretch with 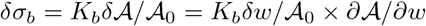. Note that provided that the basement membrane is under moderate elastic stretch before the perturbation, i.e. stretch smaller than 100%, we have that 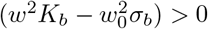.

We conclude that according to our model (see Eqn. (4)), changes in lateral surface tension *δσ_ℓ_* or changes in hydrostatic pressure *δP_H_* in the cell are predicted to trigger proportional changes in basal tension *δσ_b_*.

### E. Basal tension is increased by optogenetic activation of lateral actomyosin contractility

Utilizing our cell mechanical model described by the cellular energy functional (1), we had derived that basal tension can be increased through an increase of lateral cell tension *σ_ℓ_*, see Eq. (4). To test this prediction experimentally, we augmented lateral tension by optogenetic recruitment of RhoGEF2, an activator of actomyosin contractility, to the lateral surfaces of wing disc pouch cells, see Fig. 5A-E^33^. Optogenetic recruitment was based on the light-dependent binding of Cryptochrome 2 (Cry2) fused to RhoGEF2 (RhoGEF2-Cry2) to the N-terminal region of CIB protein (NCIB), which was targeted by a CaaX-anchor to the plasma membrane see^34,35^. We previosuly showed using this optogenetic activation that recruitment of RhoGEF2 to the lateral plasma membrane of wing disc cells results in accumulation of F-actin, increased lateral tension and decreased apical-basal cell height^35^.

**FIG. 5.**
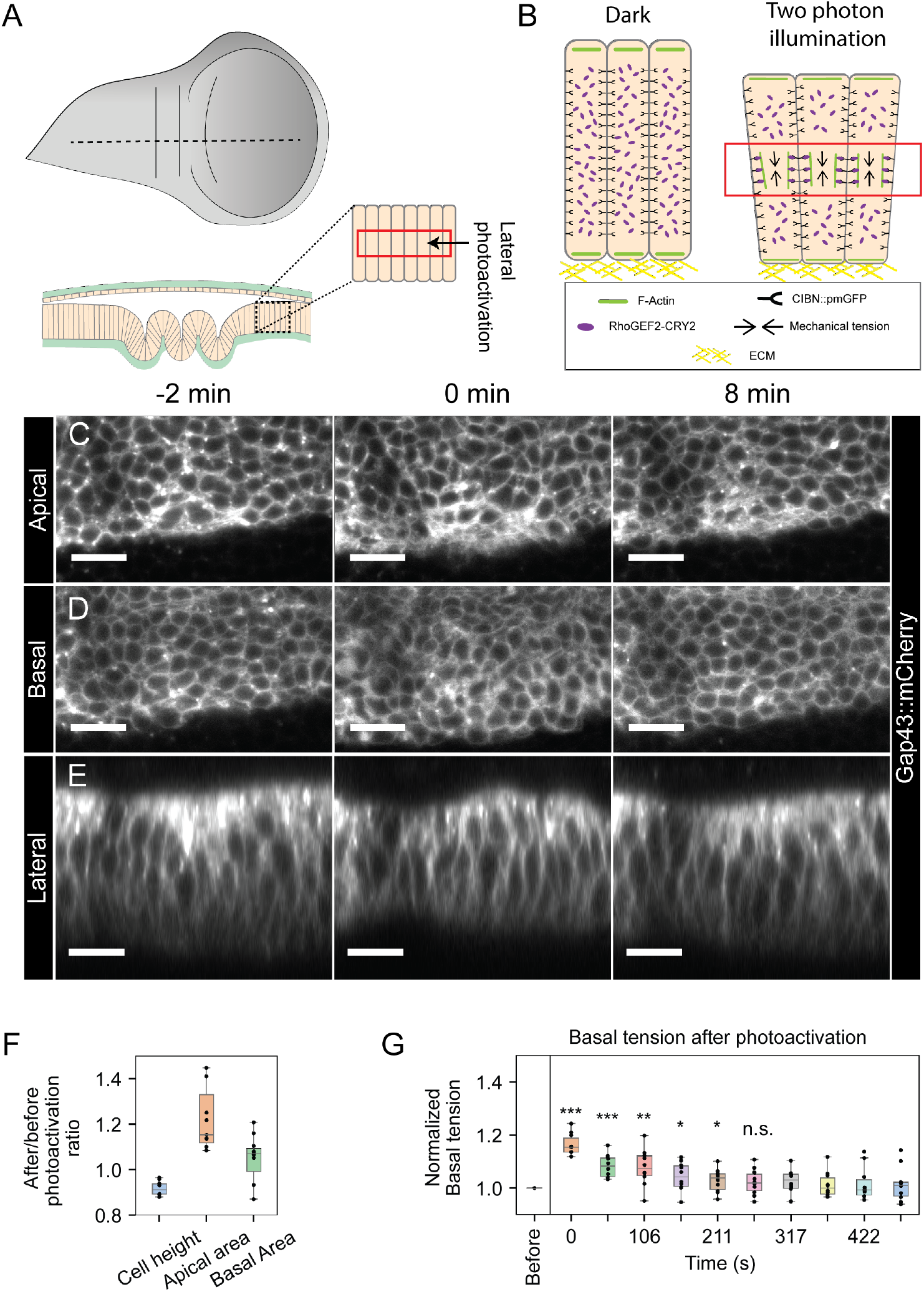
Basal tension is increased by optogenetic activation of lateral actomyosin contractility. A) Schematic illustrating the region of the wing disc that was illuminated for lateral photoactivation. The red area indicates the cross-sectional view of 15 planes, 1 *μ*m apart, that were continuously illuminated for 2 min with light of wavelength λ = 950 nm. B) Cartoon illustrating the distribution of the Rho1 activator RhoGEF2-CRY2 (purple) before and after 2 min of two-photon excitation. In the dark condition, RhoGEF2-CRY2 is localized in the cytoplasm (left). Upon two-photon excitation, RhoGEF2-CRY2 binds to CIBN::pmGFP, which localizes to the plasma membrane (right). Once at the plasma membrane, RhoGEF2-CRY2 leads to a local accumulation of F-actin (green) driving an increase in lateral tension and a reduction of cell height. (C-E) Apical (C) and basal (D) top views and lateral cross-sections (E) of a wing disc expressing RhoGEF2-CRY2 and CIBN::pmGFP, before and at the indicated times after a 2 min photoactivation. Cell membranes are labeled by Gap43::mCherry. Scale bar: 10 *μ*m. F) Boxplots of photoactivation-induced changes of cell height as well as apical and basal cross-sectional cell area (n=10 wing discs). G) Normalized basal tension before and at the indicated times after a 2 min photoactivation of RhoGEF2. Basal tension was normalized to the value before photoactivation (n=10 wing discs, same wing discs as in panel F). Significance was determined with a paired T-test (\**p* < 0.05, * * *p* < 0.01, * * \**p* < 0.001, n.s. not significant).

We illuminated a ‘lateral’ volume of wing disc pouch cells with a two photon laser of wavelength λ = 950 nm for 2 min, see Fig. 5A,B. Right after the end of photoactivation, epithelial cell height was decreased by ≈ 9%, indicating that lateral tension was increased, see Fig. 5E,F. Concomitant with the decrease in cell height, both apical and basal cell surface areas were increased by *ε_a_* ≈ 15.4% and *ε_b_* ≈ 7%, respectively, see Fig. 5C,D,F. Over time, cell shapes relaxed back to their original shape with a complete relaxation after ≈ 10 min, consistent with our previously reported observations^35^. To test whether the increase in lateral cell tension influences basal tension, we again illuminated cells with light of the wavelength *λ* = 950 nm for 2 min and used AFM indentation to measure basal tension at different time points after the end of illumination. Basal tension was increased by 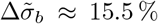 right after optogenetic activation and thereafter declined within ≈ 4 min to the level before activation see Fig. 5G. Thus, as predicted by our model, basal tension can be increased through an increase in lateral tension.

Taken together, our data demonstrate that an increase of actomyosin-based lateral cell surface tension leads to a temporary dilation of apical and basal cell surfaces with a corresponding increase of basal tension. Anticipating that the increase of basal tension is due to an increase of stretch-induced elastic stresses, we estimated the elastic modulus of the basement membrane as 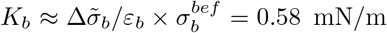 where we have used the median value of absolute basal tension before photoactivation 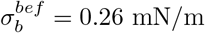.

### F. Basal tension positively correlates with hydrostatic pressure

Our cell mechanical model further predicts that a change of intracellular hydrostatic pressure *P_H_* positively correlates with basal tension, see Eq. (4). In a second set of experiments, we therefore applied mild osmotic shocks to wing discs 72 h AEL to induce a temporary change in intracellular hydrostatic pressure^36^.

To increase intracellular hydrostatic pressure, we applied a hypo-osmotic shock to explanted wing discs by diluting the culture medium down by 25% through addition of distilled water, see Materials and Methods^36^. In response, wing disc area rapidly increased by *ε* ≈ 4%, showing that the applied hypo-osmotic shock was efficient. Wing disc area subsequently relaxed to a value below the original area over a time span of ≈ 10 min, see Fig. 6A, C, perhaps through active volume control of the cells^37,38^. Concomitantly, we performed AFM measurements of basal tension. We found that right after application of the osmotic shock (measured ≈ 2 min after addition of distilled water), basal tension was increased by 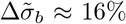 as compared to measurements before the shock. After this initial peak, basal tension relaxed to a value below the original tension, see Fig. 6D.

**FIG. 6.**
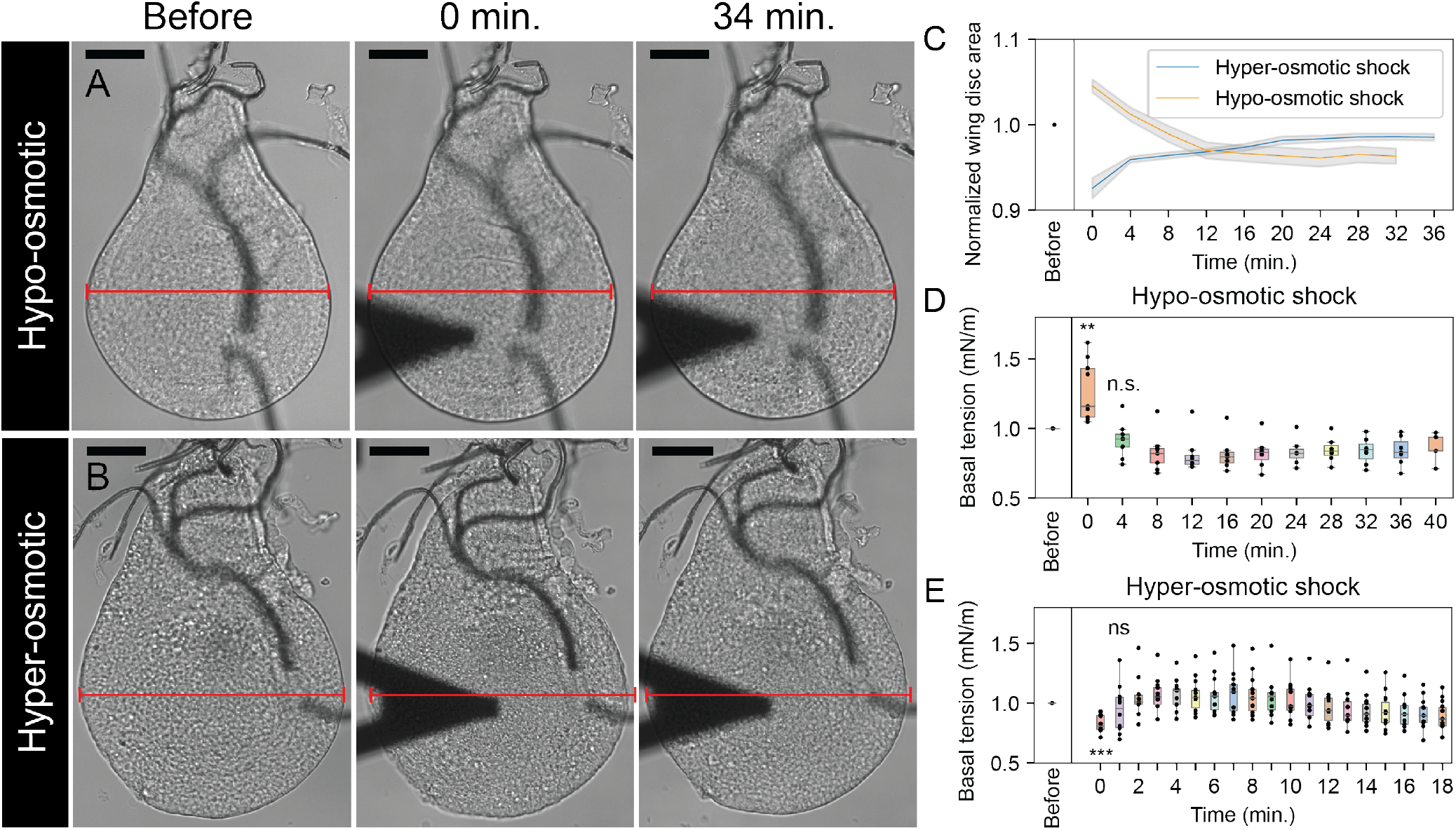
Basal tension positively correlates with hydrostatic pressure. (A-B) Brightfield time lapse images before and at the indicated times after hypo-osmotic (A) or hyper-osmotic shock (B) Scale bar: 40 *μ*m. The red line is of constant length and indicates changes of horizontal wing disc dimension with the left line edge coinciding with the left wing disc edge. C) Normalized wing disc area before and at the indicated times after hypo-osmotic (orange) or hyper-osmotic (blue) shock (n=10 wing discs). The areas were normalized by the value before the osmotic shock. Mean and s.e.m. are shown. (D-E) Boxplots of basal tension before and at the indicated times after hypo-osmotic (D) or hyper-osmotic (E) shock. The tension was normalized by the value before the osmotic shock (n=10, same wing discs as in panel C). Significance was determined with a paired T-test (*\*p* < 0.05, * * *p* < 0.01, * * *\*p* < 0.001) n.s. not significant

To decrease intracellular hydrostatic pressure, we applied a hyper-osmotic shock to explanted wing discs by increasing the concentration of the culture medium through addition of sorbitol to a final concentration of 13 mM, see Materials and Methods^36^. In response, wing disc area was decreasing by ≈ 6% with a subsequent recovery to its original value over a time span of ≈ 20 min, see Fig. 6B, C. Accompanying AFM measurements showed a reduction in basal tension by ≈ 19% right after application of the hyper-osmotic shock (measured ≈ 2 min after addition of sorbitol). Subsequent to this initial drop, basal tension recovered to its original value, see Fig. 6E. Thus, as predicted from our model, changes in hydrostatic pressure positively correlate with changes in basal tension (and wing disc area) during, both, hypo-osmotic and hyper-osmotic shock.

Again based on the assumption that basal tension changes upon osmotic shock were mainly triggered through changes in elastic stretch of the basement membrane, we used our experimental results to estimate the elastic modulus of the basement membrane *K_b_*. For hypo-osmotic shocks, median tension values before osmotic shock were 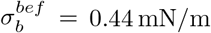. Together with a relative tension increase of 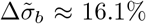 and a surface area increase of *ε* ≈ 4.1%, we can estimate 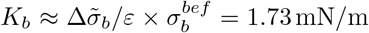. On the other hand, for hyper-osmotic shocks, median basal tension values before osmotic shock were 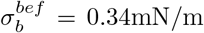. Together with a relative tension decrease of ≈ 19.1% and a surface area decrease of ≈ 6.1%, we estimated *K_b_* ≈ 1.06 mN/m. We speculate that the larger elastic modulus for hypo-osmotic stretching might originate, at least partly, in the nonlinear strain-stiffening of basement membranes as has been reported previously^39,40^.

## III. DISCUSSION

In this study, we quantified basal tension in the columnar epithelium of explanted wing discs from the 3rd instar larva of the fruit fly *Drosophila melanogaster*. We used an established method of AFM indentation to estimate basal cell surface tension by the scaled slope of force-indentation curves, see Fig. 1 and Fig. S1^20,21^. In particular, we find that basal tensions in wing disc epithelia are in the range of 0.2 – 0.5 mN/m and therefore similar to tension values of the actin cortex of isolated interphase cells measured with diverse methods^22–25^. Previous estimates of basal tension in epithelia have been reported; micropipette aspiration experiments on mouse blastocysts and 8-cell stage mouse embryos provided similar basal tension estimates between 0.5 – 1 mN/m and 0.2 – 0.5 mN/m, respectively^41,42^. By contrast, Yang *et al*. determined basal tensions in intestinal organoids by micropipette aspiration finding considerably higher values between 3 – 6 mN/m^43^. Taken together, these results suggest that basal tension in epithelia may vary by at least one order of magnitude perhaps in dependence of hydrostatic pressure in the lumen.

Perturbing wing discs pharmacologically, we find that both basement membrane and actomyosin contribute to basal tension generation albeit with a bigger influence of the basement membrane, see Fig. 1D-F. In accordance with our study, Yang et al. and Maitre *et al*. found evidence that actomyosin contributes to basal tension generation^42,43^. Furthermore, ablation experiments on basal cell edges previously showed that basal tension is reduced by basement membrane degradation through incubation with Collagenase or localized MMP2 expression^19^.

In order to understand how the basement membrane contributes to basal tension, we investigated the material properties of the basement membrane and in particular asked whether it behaves solid- or liquid-like on the time scale of fold formation. Performing imaging of Collagen IV in the basement membrane for several hours, we observed that turnover and mobility of Collagen IV is slow (≈ 40 h) suggesting a solid-like nature of the basement membrane on time scales of several hours, see Fig. 3. This observation of slow basement membrane turnover is in accordance with the majority of earlier reports about the material properties of basement membranes^1^. The basement membrane turnover in *Drosophila* embryos was, however, measured to be slightly faster (7-10 h)^44^.

Given that the basement membrane has solid-like properties, we conclude that the basement membrane can be stretched elastically and store elastic stresses on long time scales thereby contributing to basal tension. The presence of elastic stretch in the basement membrane of the wing disc epithelium has also been reported by Harmansa *et al*. who linked ECM-associated elastic stresses to epithelial thickening and doming in the wing disc^45^.

Passive elastic stretch and corresponding elastic stresses in the basement membrane need to be maintained by active cellular driving forces. As potential candidates, we addressed actomyosin-dependent contractility in lateral cell faces and intracellular hydrostatic pressure. To test these two potential driving forces, we perturbed the wing disc epithelium through i) optogenetic actomyosin activation in lateral cell faces and ii) hypo-osmotic wing disc swelling. In response, we found that basal cell areas were stretched and basal tension increased, see Fig. 5 and Fig. 6. In a complementary experiment, we performed hyper-osmotic wing disc shrinkage observing a concomitant reduction in basal cell areas and basal tension, see Fig. 6. We conclude that our experimental results are in support of the hypothesis that lateral actomyosin contractility and hydrostatic pressure drive basement membrane stretch.

We note that the scenario of an elastic basement membrane stretch balanced by an active expansile cell layer is further supported by our observation of basal cell area widening upon Collagenase-induced basement membrane degradation, see Fig. 1I. Furthermore, basal area shrinkage upon treatment with the Rock inhibitor Y27632 and Latrunculin A, see Fig. 1J,K, supports that basement membrane stretch is maintained by actomyosin forces.

From measurements of basal tension changes during induced wing disc area changes, we could infer estimates of the basal elastic modulus in the range of 0.5 – 2mN/m. Using previously reported thickness estimates of 0.1 *μm* of the wing basement membrane in the 3rd instar larva^46^, we can associate a basement membrane Young’s modulus of ≈ 5 – 20 kPa assuming a Poisson ratio of 0.5^47,48^. This estimate is in the lower range of previously measured elastic moduli of basement membranes which were reported to lie between 1 – 1000 kPa.

In summary, data presented here suggest that basal tension in wing disc epithelia is to a large extent regulated through elastic stretch in the basement membrane and corresponding elastic stresses in the basement membrane material. Importantly, this indicates that basal tension generation has quite distinct properties from apical tension generation which relies on actomyosin contractility; the actin cortex turns over on time scales of few tens of seconds and, thus, cannot provide a shape memory on time-scales longer than its turnover time^22,49–51^. Therefore, corresponding tension is mainly set by the momentary molecular configuration of the actin cortex. By contrast, the basement membrane ECM is a more solid-like material than the actin cortex and possesses a shape memory on the time scale of hours and longer^1^. Correspondingly, basal tension is set, to a large extent, by the deviation from the resting basal area in conjunction with the stiffness and thickness of the basal ECM network. We suggest that the asymmetry in apical and basal tension generation is an important tool of morphogenesis to induce asymmetries in apical and basal tension, which can then lead to curvature in the epithelium.

## IV. MATERIALS AND METHODS

### A. Fly stocks and genetics

Flies were raised on standard fly food and mantained at 25°C unless stated otherwise. The following *Drosophila melanogaster* fly stocks were used:

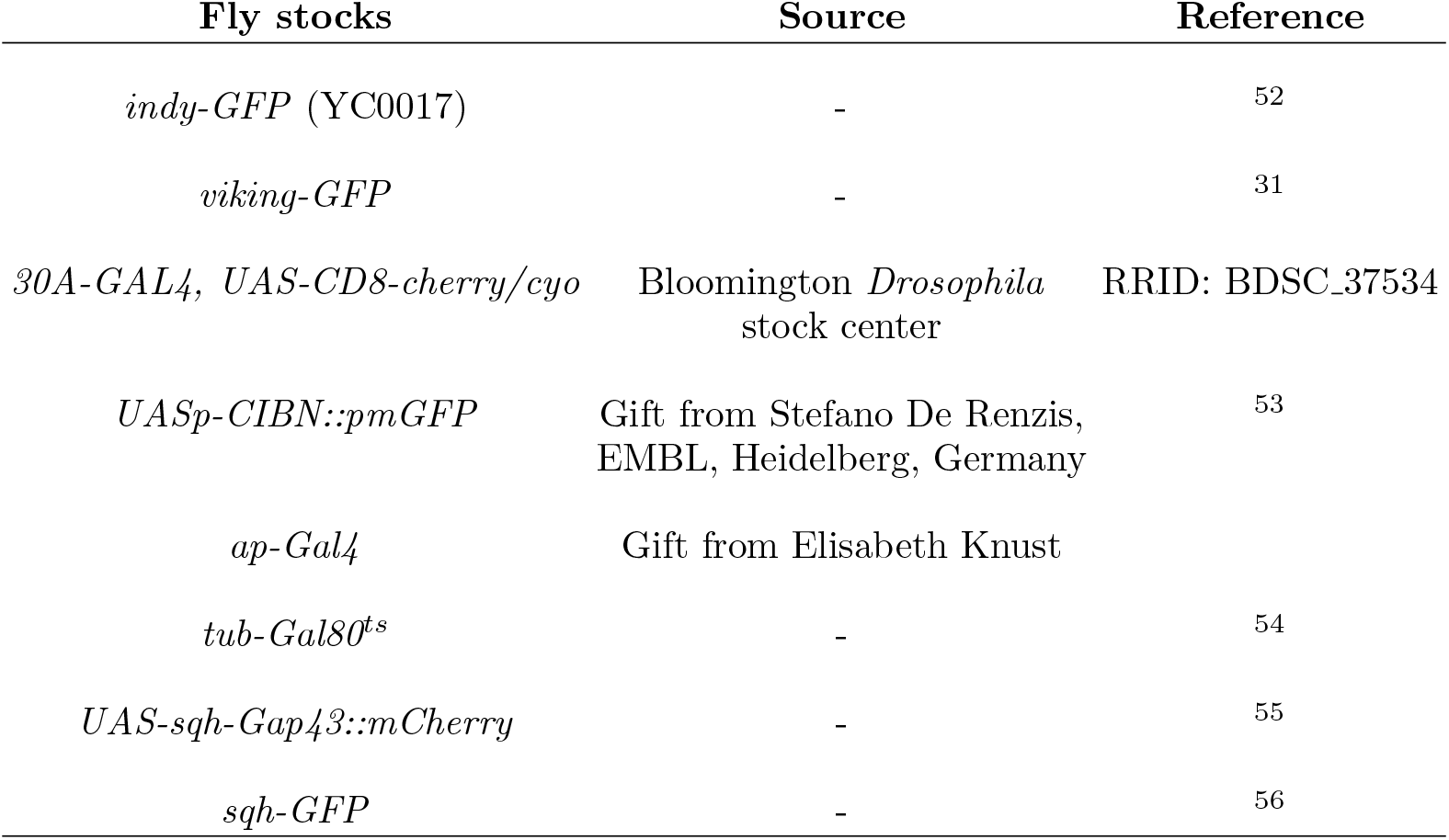

Standard fly husbandry and genetic methodologies were used to cross *Drosophila* strains. The detailed genotype for each experiment were as follows:

| Figure 1 | Genotype |
| --- | --- |
| B,D-G | *y w;vkg-GFP,30A-GAL4, UAS-CD8-cherry/cyo* |
| H | *sqh-GFP* |
| I-K | *indy-GFP* |
| Figure 2 |  |
| B | *y w;vkg-GFP,30A-GAL4, UAS-CD8-cherry/cyo* |
| C-F | *indy-GFP;30A-GAL4, UAS-CD8-cherry/cyo* |
| Figure 3 |  |
| A-I | *y w;vkg-GFP* |
| Figure 5 |  |
| A-G | *ap-Gal4, tub-Gal80ts, UASp-CIBN::pmGFP; UASp-RhoGEF2-CRY2/sqh-Gap43::mCherry* |
| Figure 6 |  |
| A-E | *y w;vkg-GFP,30A-GAL4, UAS-CD8-cherry/cyo* |
| Supplementary Figure 1 |  |
| A-C | *y w;vkg-GFP,30A-GAL4, UAS-CD8-cherry/cyo* |
| D-F | *sqh-GFP* |
| G-I | *indy-GFP* |

### B. Time lapse live imaging and FRAP experiments

Staging, dissection and culture of 72 h AEL wing discs were done as described previously^19^. Imaging as reported in Fig. 3 was performed using a Zeiss LSM700 confocal microscope of the CMCB light microscopy facility, using a 40x/1.25 numerical aperture water immersion objective. Image stacks of 30 slices were taken such that the total height of the wing disc was covered (≈ 40 *μ*m). The pinhole aperture was chosen such that the optical section thickness was equal to the distance between neighboring slices. Wing discs were mounted with their basal side towards the objective. We chose a pixel number of 512 × 512 pixels per image and recorded with 2% laser power.

To observe Collagen IV dynamics long-term (Fig. 3A-E), we carried out the following imaging protocol: First, an initial z-stack of the whole tissue was acquired. Then, two square-shaped regions of interest (30 × 30 *μ*m) were chosen, one in the hinge/notum region and another one in the pouch. Subsequently, we imaged a time series of 42 frames of both regions with a time interval of 10 min for a total time of ≈ 7 h. In the end, a final z-stack of the whole wing disc was acquired.

Fluorescence recovery after photobleaching (FRAP) experiments as described in Fig. 3F-I were performed in a similar manner; first, an initial z-stack of the full wing disc was recorded. Then, photobleaching was performed on a square region (30 × 30 *μ*m) with a 488 nm laser at 100% power for 28 seconds giving approximately 90% fluorescence intensity reduction. Subsequently, a times series of 42 z-stacks of a 45 × 45 *μ*m square region (bleached region with surroundings) were recorded with a time interval of 10 min. In the end, a final z-stack of the full wing disc was acquired. All image analysis was performed using FIJI^57^ as described for the confocal imaging. All measurements were performed at room temperature.

For both, long-term imaging with and without photobleaching, we performed image analysis using FIJI^57^ to measure the mean of the summed stack intensities of square regions of interest over time (Fig. 3D,H). Fluorescence intensities were normalized by the fluorescence intensity value at the first time point.

To quantify the influence of bleaching introduced through long term imaging (Fig. 3A-E), the ratio of initial and final mean fluorescence intensity was determined for both square regions and compared to an analogous fluorescence intensity ratio for a square region of equal size outside of the long-term imaged square regions (Figure 3C). This region is labeled as ‘OR’ and depicted with a cyan frame, see (Figure 3A,C).

For the fluorescence line profile in Fig. 3I, we chose a line thickness of approximately one third of the bleaching frame height.

### C. Acquisition and analysis of AFM force-indentation curves

Wing disc indentations were performed using an atomic force microscope (Nanowizard I, JPK Instruments) mounted on a Zeiss Axiovert 200M wide-field microscope, 20x objective (Zeiss, Plan Apochromat, NA=0.8) along with a CCD camera (DMK 23U445 from Theimagingsource).

During measurements, dissected wings discs were placed into cell culture dishes (fluorodish FD35-100) coated with poly-D-lysine. The coating was performed by adding 2 ml of poly-D-lysine solution (0.1 mg/ml in PBS, Gibco A3890401) to the dishes and incubating over night at 4°C. Then, the poly-D-lysine solution was removed and dishes were rinsed once with distilled water. Afterwards, we dried the plates with a compressed air gun. Plates were stored at 4°C up to five days.

AFM cantilevers with pyramidal tips were used in all experiments with a nominal spring constant of 0.01 N/m (MLCT-C, Bruker AFM probes). Preceding each experiment, the actual cantilever spring constant was measured by means of thermal noise analysis (built-in software, JPK). Actual spring constant values ranged from 0.01–0.02 N/m. All measurements were performed in 3 ml of Grace’s culture medium at room temperature. AFM measurements were carried out in contact mode. For tension measurements presented in Fig. 1, 2B, 6D, a measurement consisted of 64 force indentation curves acquired on a regular square lattice of 8 × 8 points with an edge length of 10*μ*m. For tension measurements presented in Fig. 5G and 6E, a measurement consisted of 16 force indentation curves acquired on a regular square lattice of 4 × 4 points with an edge length of 10 *μ*m.

For all grid configurations, indentations were carried out continuously such that the cantilever was indenting one grid point after another. After completion of the full grid, the indentation protocol started over. Indentations were performed at an extension speed of 1 *μ*m/s up to a maximal force of 500 pN. The position of the grid was targeted by eye in the center of the wing pouch unless stated otherwise.

For position-dependent measurements shown in Fig. 2C-F, force indentations were performed on a rectangular grid with 8 × 14 grid points spanning an area of 25 × 65 *μ*m^2^ (68 h AEL) or 6 × 20 grid points spanning an area of 25 × 100 *μ*m^2^.

In order to get the slope of the force-indentation curves, we used the JPK Data Processing software. First we applied a baseline subtraction, including the correction for the offset plus tilt of the curves. This was followed by a subtraction of the *x*-offset (contact point) and a correction for the bending of the cantilever (vertical tip position). Afterwards, we performed a linear fit in the range between the contact point and 500 pN. The data processing and plotting was performed using Python.

The median of the successfully fitted ≈ 64 curves represents the basal tension of one sample and corresponds to one data point in boxplots in Figs. 1D-G, 2B, 5G, 6D,E and Supplementary Figs. S1 and S2D,H.

For tension values on an X-Y grid as reported in Fig. 2 C’,E’, indentations were performed three times per grid point in every wing disc. The tension value of each grid point is the average of three indentation measurements. To generate the curves in Fig. 2D,F, we calculated the median of tension values measured along the Y-axis for a fixed X position for each wing disc. Tensions were then normalized by the tension value in the center of the hinge region (*X* = 0). Finally, normalized basal tension were averaged over several wing discs and plotted across the X position (number of wing discs averaged: n=7 for the 68 h AEL and n=11 for the 72 h AEL wing disc).

### D. Wing disc drug treatment and AFM measurements

*Ex vivo* drug treatment was applied to wing discs explanted 72 h AEL. For that purpose, wing discs were transferred to a coated glass bottom dish and immersed in 3 ml culture medium as explained in Section IV C. Then, an initial measurement series of AFM indentations along an 8 × 8 grid was performed in the center of the pouch region. Afterwards, the AFM head was unmounted and the respective drug was added to the medium. To ensure proper mixing, pipette pumping was performed. Afterwards, the AFM head was mounted again as quickly as possible (~ 2 min) and AFM indentations along an 8 × 8 grid were continuously carried out for approximately 30 min (see Fig. 1D-G and Supplementary Fig. S1A-C). AFM force-indentation curves were analyzed to obtain basal tension estimates as described in Section IV C. The time evolution of basal tensions after drug addition are shown in Supplementary Fig. S1A-C. For all drug treatments, we concluded that the full effect on wing disc mechanics was present at the latest after 15 min of treatment time.

The following drug concentrations were used: 1 mM Y27632 (Biomol AG-CR1-3564-M010, 25 mM stock in PBS), 0.2 % Collagenase Type I (Sigma-Aldrich SCR103, 1% stock in PBS) 4 μM Latrunculin A (Biomol AG-CN2-0027-C100, 1 mM stock in DMSO). In a control measurement, AFM indentations were performed before and after treatment with DMSO at final concentration of 2 % in the culture medium (see Fig. 1D, rightmost panel).

For measuring the change in apical and basal area after drug treatment, cell membranes in the pouch region were visualized by Indy-GFP. Wing imaginal discs were mounted with their apical side tethered to the glass bottom dish. Imaging was performed using a Leica SP8 MP confocal microscope. Image analysis of drug treatment was done using FIJI^57^. In order to be able to visualize the basal surface of the cells, the signal-to-noise ratio was improved using a machine-learning image restoration plugin (Content-Aware Image restoration, CARE^58^). The apical plane was identified by the accumulation of signal intensity. The basal plane was identified focusing the image plane on the basal surface of cells. Groups of cells (5-6 cells per group) were then segmented and tracked over time using Tissue Analyzer^59^. The area of the group was measured using Tissue Analyzer^59^ or Fiji^57^.

To quantify Myosin levels at the basal surface of the cells, Myosin acitivity was visualized by Sqh-GFP. Imaging was performed using an upright Leica Stellaris 8 confocal microscope. Wing imaginal discs were mounted with their apical side tethered to the glass bottom dish. Image Z-stacks were taken from basal to apical every 2 min. Myosin intensity was measured on a squared region (30 × 30 *μ*m) using Fiji^57^. The peak of the mean fluorescence intensity z-profile was used to identify the basal surface.

### E. Optogenetics experiments and analysis

Incubation, dissection and culture of wing discs were done as described previously^19,35^. Larvae were incubated in the dark at 25°C and transferred to 29°C 3 days before dissection. Wing discs were placed into cell culture dishes (fluorodish FD35-100) coated with poly-D-lysine. Two-photon activation and imaging were performed via a Zeiss LSM 980 confocal laser scanning microscope, using a C-Apochromat 40x/1.2 water objective.

For every wing disc measurement, the following protocol was carried out: First, wing discs were mounted with the apical side facing the objective. A region of interest (ROI) of 50 × 30 *μ*m was defined in the central pouch region of wing discs. Before photoactivation and AFM measurements, an initial z-stack in the mCherry channel (561 nm) was acquired (45 planes, 1 *μ*m spacing). Two-photon photoactivation was performed with light of 950 nm wavelength at 8% laser power with bidirectional scanning. A 15 *μ*m thick z-interval in the center of the wing disc epithelium was photoactivated for 2 min by scanning 15 planes 1 *μ*m apart. After photoactivation, a z-stack (45 planes, 1 *μ*m spacing) of the region of interest was recorded in the mCherry channel in a time series (5 frames, 2 min interval). After verifying the effectiveness of the optogenetic activation by imaging, an initial AFM measurement was acquired on a regular square lattice of 4 × 4 on the tissue, see Section IVC above. Afterwards, a second 2 min lateral photoactivation was carried out immediately followed by continuous AFM indentations on a 4 × 4 grid for 10 minutes (without confocal imaging).

Image analysis of optogenetic experiments was carried out in FIJI^57^. To quantify cell height in the wing disc epithelium, a vertical cross-section was acquired using the reslice tool on a middle section of the imaged region. In the cross-section, we acquired the fluorescence intensity profile on a line (≈ 30 *μ*m width) encompassing the full z-range. We subtracted the background fluorescence intensity and quantified tissue height as the length of the line interval where the fluorescence intensity was reduced by no less than 20 % as compared to the maximal fluorescence intensity value. Apical cell areas were determined by choosing a plane between the first 5 to 10 slices. Afterwards, we measured the area of a set of cells (4-5) whose apical region stayed focused in the same plane before and after photoactivation. The procedure for measuring basal cell areas was similar to apical cell areas. The plane was chosen between the last 5 to 10 slices of the z-stack.

### F. Osmotic shocks

Osmotic shock experiments in conjunction with AFM measurements were carried out analogous to AFM measurements before and after reagent treatment as described in Section IV D. Instead of reagent additions, 1 ml of dH_2_0 was added to 3 ml of medium in the dish to reduce the medium osmolarity to 75 % for hypo-osmotic shock experiments. For hyper-osmotic shock experiments, sorbitol (Sigma S1000000) solution was added to a final concentration of 13 mM to the medium.

### G. Scaled force indentation slopes as a quantitative estimate of basal tension

In the search for a mechanical model for indentation analysis, we noted that corresponding force-indentation curves of untreated wing discs appeared as a roughly linear force increase *F* ∝ *δ*, i.e. a power law increase with exponent one. The Hertz model of indentation into an elastic half space, predicts a parabolic increase of the force *F* ∝ *δ*^2^ corresponding to a power law with exponent two^20,21,60^. Upon closer inspection, we found that fitting power laws with arbitrary exponents to measured force-indentation curves yields on average exponents of ≈ 1.35, see Fig. S1. We therefore concluded that standard analysis with the mechanical model of indenting into an elastic half space is not an appropriate analysis method for our data.

Motivated by the scenario that the basement membrane might be the dominant structure in mechanical response, we turned to the mechanical model of indentation into a mechanically tensed elastic sheet. In fact, theoretical studies predict a power law exponent one for shallow indentation into a tensed, disk-shaped elastic sheet which is clamped at the periphery^61,62^. Therefore, the power law predicted for this scenario matches more closely the force increase observed in our experiments.

Our hypothesis that the basement membrane is the dominant mechanical structure is further supported by indentation experiments before and after wing disc treatment with Collagenase. Treatment of wing discs with Collagenase leads to a disintegration of the basement membrane. Upon treatment, we see a significant softening of the wing disc over time, see Fig. S1A. Furthermore, softening is accompanied by an increase of the power law exponent to a value of ≈ 2, see Supplementary Fig. S2B,C, which matches closely the power law predicted for indentation into an elastic half space. This observation suggests that only after basement membrane removal, the cytoplasmic bulk acts as the dominant mechanical element in AFM indentation.

Taken together, these observations motivated us, to analyse obtained AFM data by applying the mechanical model of indentation into a thin, tensed elastic sheet which we identify with the composite layer constituted by the basement membrane-actin cortex sandwich at the basal side of the columnar epithelium. Provided that tension is sufficiently large, theoretical studies predict *F* = *πσ_b_δ* for indentations of a clamped, tensed elastic sheet to which a central point force is applied. Here *σ_b_* is the tension due to elastic membrane prestretch and δ is the cantilever indentation^61,62^. Correspondingly, we estimate basal tension in the wing disc as 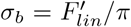 throughout this manuscript, where 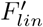 is the fitted linear slope of the measured force-indentation curve, see Fig. 1C. We note that this analysis scheme has been previously used to determine actin cortical tension in single cells and epithelial sheets^20,21^.

### H. Power law fitting of force-indentation curves

The matlab code for power law fitting of force-indentation curves is provided as part of the Supplementary material (“FitAndAverageForceCurves.m”). In short, force curves corresponding to the measurements on one wing disc at a specific time point are read in by the program and an average force indentation curve is calculated. By least squares fitting, the contact point of the averaged force-indentation curve is readjusted such that the curve is best captured by a linear increase in the log-log representation of the data (i.e. by a power law). Then, a second least squares fit is performed to determine the optimal power law exponent. For untreated wing discs (dissected 72 h AEL), the power law exponent is typically between 1.2 – 1.4 (see Supplementary Fig. S2) and therefore closer to 1 than to 2.

## Supporting information

Supplementary script

## ACKNOWLEDGMENTS

We thank Frank Jülicher and Carsten Werner for fruitful discussions on the topic. Furthermore, we thank Marko Brankatschk for giving generous access to his dissection microscopes and fly incubators. CD and EFF acknowledge financial support from the Deutsche Forschungsgemeinschaft under Germany’s Excellence Strategy, EXC-2068-390729961, Cluster of Excellence Physics of Life of TU Dresden. In addition, the authors thank the CMCB Light Microscopy Facility for excellent support.

