## Supplementary for "Hydrostatic pressure and lateral actomyosin tension control stretch and tension of the basement membrane in epithelia"

K.Y. Guerra Santillán,[1,2] C. Dahmann,[2,1,a)] and E. Fischer-Friedrich[1,3,2,b)]
[1)] *Cluster of Excellence Physics of Life, Technische Universität Dresden, Dresden, Germany*
[2)] *School of Science, Technische Universität Dresden, Dresden, Germany*
[3)] *Biotechnology Center, Technische Universität Dresden, Dresden, Germany*

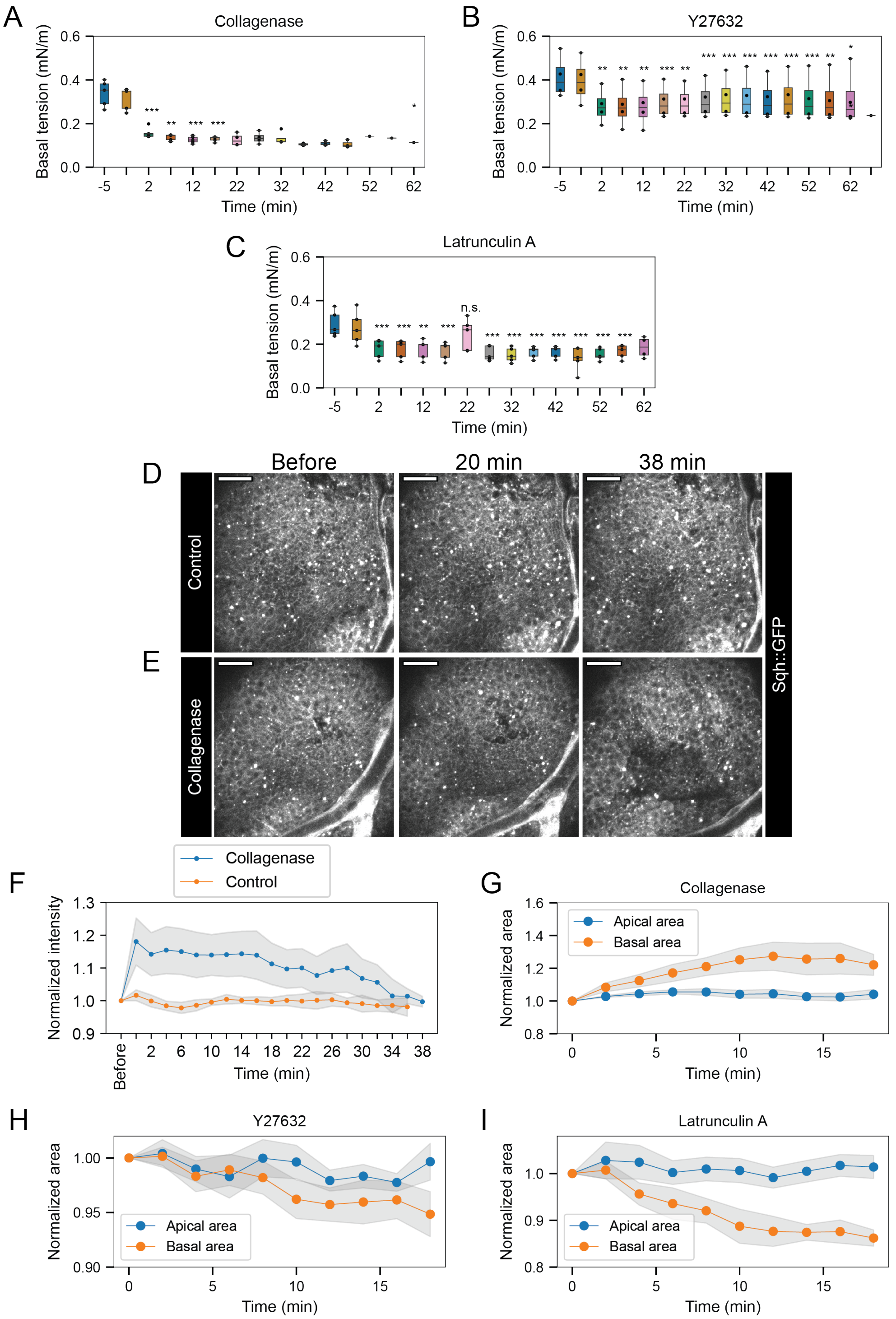


FIG. S1. Time evolution of basal tension in wing discs in response to treatments with reagents A) Collagenase (0.2 mg/ml), B) Y27632 (1 mM) and C) Latrunculin A (4 $\mu$M) for the indicated times ($N = 4$, each). Negative times indicate that the reagents had not yet been added to the medium. Data correspond to boxplots in Fig. 1D-F, main text. Significance was determined with a paired T-test (*p $<$ 0.05, **p $<$ 0.01, ***p $<$ 0.001, n.s. p$>$ 0.05 not significant). (D-E) Exemplary sum fluorescence intensity projections of a Z-stack of images of the basal part of a 72 h AEL wing disc expressing Sqh-GFP, before and at the indicated times after incubation in culture medium (D) and in medium with Collagenase (E). The concentration is the same as in A). Scale bars: 20 $\mu$m. F) Mean and s.e.m. normalized Sqh-GFP fluorescence intensity along a $30 \times 30\,\mu$m square in control conditions (orange) and with Collagenase (blue) for n=4 and n=5 wing discs, respectively. The fluorescence intensity was normalized by the initial value of each wing disc. Data correspond to boxplots in Fig. 1H. (G-I) Mean changes of apical (blue) and basal (orange) cell areas at the indicated times (Collagenase, Latrunculin: N=5, Y27632: N=7, same concentrations as in Fig. 1D-F). The grey area indicates the standard error of the mean. Data correspond to boxplots in Fig. 1I-K.

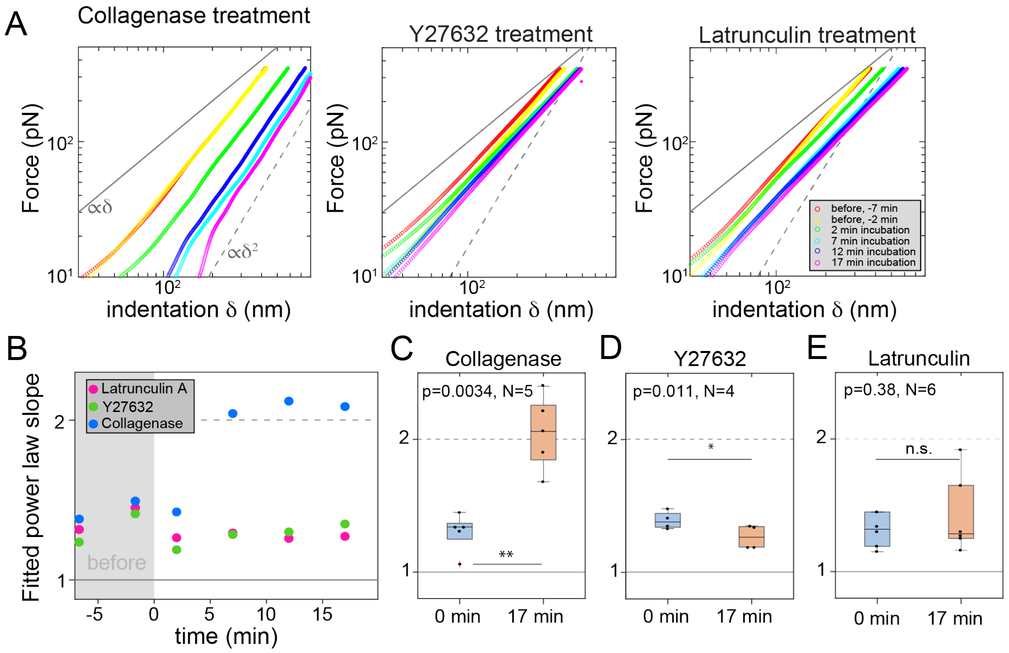


FIG. S2. Fitted power law slope of force-indentation curves obtained by AFM indentation with a pyramidal indenter on the basal side of the pouch of a wing disc dissected at 72 h AEL. The mechanics of each wing disc was first probed in regular medium (control). After this, the reagent indicated was added to the medium. The evolution of wing disc mechanics during reagent incubation was measured over time. A) Log-log-plot of force indentation curves before reagent addition (red and yellow) and for growing incubation times with indicated reagents (green: 2 min incubation, cyan: 7 min incubation, blue: 12 min incubation, pink: 17 min incubation time) measured on one exemplary wing disc. Each curve represents an average over 64 indentation points. The grey lines indicate a power law growth with exponent 1 (solid grey) or exponent 2 (dashed grey). Theoretical studies predict a power law exponent 1 for shallow indentation into a prestressed elastic sheet[1–3] and a power law exponent 2 for indentation into a solid elastic half space[3–5]. Left panel: treatment with Collagenase (0.2 mg/ml), middle panel: treatment with Y27632 (1 mM), treatment with Latrunculin A (4 $\mu$M). Each colored curve represents the average of force-indentation curves measured on a $8 \times 8$ grid with edge length 10 $\mu$m (see Materials and Methods). B) Time evolution of fitted power law exponent of averaged force-indentation curves before addition of the reagent (grey area) and after the indicated incubation time for representative wing disc (same data as in panel A). While there is no meaningful change of the exponent for treatment with Y27632 and Latrunculin A, incubation with Collagenase significantly increases the power-law exponent. C-E) Boxplots displaying fitted power law exponents before wing disc treatment and after 17 min reagent incubation: C) Collagenase treatment (wing discs measured: N=5), D) Y27632 treatment (wing discs measured: N=4), E) Latrunculin A treatment (wing discs measured: N=6). Significance was determined with a paired T-test (*$p < 0.05$, **$p < 0.01$, ***$p < 0.001$).
