## Supplementary script for "Hydrostatic pressure and lateral actomyosin tension control stretch and tension of the basement membrane in epithelia": FitAndAverageForceCurves.m

```
function AverageScriptForce()
close all
times=[-5, 0, 0, 5, 10, 15, 20, 25];
figure(2);hold off; figure(1); hold off; figure(3); hold off;

%%measurement of wing disc 1 with collagenase
pathst='~/mypath/ForceCurves_Drugs/WD_collagenase_19012021/';%
times=[-5, 0, 2, 7, 12, 17, 22, 27, 32, 37, 42, 47, 52];
% list of folders that contains force indentation curves measured at
% different time points
paths={[pathst '-5min_processed-2021.07.15-11.54.29/']; [pathst '0min_processed-2021.08.13-14.14.59/'];...
    [pathst '0bismin_processed-2021.08.13-14.19.12/']; [pathst '5min_processed-2021.08.13-14.17.46/'];
    [pathst '10min_processed-2021.08.13-14.12.15/']; [pathst '15min_processed-2021.07.15-12.35.52/'];...
    [pathst '20min_processed-2021.08.16-16.44.54/'];[pathst '25min_processed-2021.08.16-16.45.31/'];
    [pathst '30min_processed-2021.08.17-10.25.40/']; [pathst '35min_processed-2021.08.17-10.47.11/'];
    [pathst '40min_processed-2021.08.17-10.44.11/']; [pathst '45min_processed-2021.08.17-10.46.06/'];
    [pathst '50min_processed-2021.08.17-10.45.28/']; };
Fmax=350; 
xmin=-1200; xmax=500; %range of indentations


Colors=[[0.3 0 0.6]; [0 0 1]; [0 0.5 1];[0 1 0.5]; [0 1 0];[0.5 1 0];[0.5 0.5 0];[1 0 0]];
Colors=hsv(length(paths));
slopes=zeros(1,length(paths));
shifts=slopes;
LegendStrings=cell(length(paths),1);
newopts=optimset('lsqcurvefit');
optnew = optimset(newopts,'TolX',1e-18, 'MaxFunEvals',2800);
shift0=70; linfunc = @(p,x) p(1)*x+p(2); para0=[1 0];

for ind=1:length(paths)
    try
        temp=dir(char(paths(ind))); paths(ind)={[char(paths(ind)) temp(end).name '/']};
        files=dir([char(paths(ind)) '*.txt']);
        slopes2=zeros(1,length(files));
        shifts2=zeros(1,length(files));
        xvec=xmin:xmax; % admits to consider indentations down to -0.5 um
        ForceCurves2=zeros(length(files),length(xvec));
        for fileidx=1:length(files)
            ForceInd=importdata([char(paths(ind))  files(fileidx).name],' ',77);
            ForceIndData=ForceInd.data;
            ForceIndHeader=ForceInd.textdata(2:end,1);
            Kspring=str2num(ForceIndHeader{71}(19:25))*1000;%unit: pN/nm
            VertDefl=smooth(ForceIndData(:,2)*10^12,10); % force in pN
            temp=VertDefl;
            HeightMeas=ForceIndData(:,end-3)*10^9; % force in Nanometer
            HeightCorr=HeightMeas+0*VertDefl/Kspring; % factor 0 before VertDefl if cantilever bending was already taken into account in the analysis
            ForceCurves2(fileidx, :)=interp1(HeightCorr,VertDefl,xvec);
            yvec=ForceCurves2(fileidx, :);
        end
        
        temp=char(paths(ind,1));
        LegendStrings(ind)={[temp(length(pathst)+1:length(pathst)+6) ', N=' num2str(sum(~isnan(mean(ForceCurves2,2,'omitnan'))))] };
        
        yvec=mean(ForceCurves2,'omitnan'); % averaging of forces measured at one time point
        yvec=yvec-min(yvec); temp=yvec; yvec=yvec(temp<Fmax); xvec=xvec(temp<Fmax);
        
        [C,I]=min(yvec); 
        %xcon=xvec(min(find(yvec<min(yvec)+max(yvec)/10000)));%find rough estimate for contact point, option 1
        xcon=xvec(I)-20; %find rough estimate for contact point, option 2
        
        
        newopts=optimset('lsqcurvefit'); shift0=xcon;
        optnew = optimset(newopts,'TolX',1e-18, 'MaxFunEvals',2800);
        
        shift0=70;
        % least square fit for fine adjustment of the contact point 
        % a small shift of the rough guess xcon is calculated such that the force-indentation curve is best captured by a linear increase in the log-log representation
        [shift,resnorm,residual,exitflag,output,lambda,jacobian] =lsqnonlin(@CalcR2, ...
            shift0, [-350],[350], optnew, xvec, yvec, ind);
        shifts(ind)=shift;
        
        xvec=xvec-shift;
        figure(1); plot(xvec, yvec, 'Color', Colors(ind,:)); hold on;
        
        figure(2);
        loglog(-xvec,yvec,'o', 'MarkerEdgeColor', Colors(ind,:));
        hold on;
        ylim([10,500]); xlim([30,800])
        linfunc = @(p,x) p(1)*x+p(2); para0=[1 0];
        % least square fit, that finds the best power law exponent
        [para1,~,resid,~,~,~,jacob] = lsqcurvefit(linfunc,para0,log(-xvec(yvec>10 & xvec<0))/log(10),(log(yvec(yvec>10 & xvec<0)))/log(10));
        slopes(ind)=para1(1); % slope of force indentation curve measured at specific incubation time point
    catch
    end
end
plot(1:400, (1:400),'k', 'LineWidth',2)
plot(1:400, (1:400).^2/600,'k', 'LineWidth',2)
legend(LegendStrings);
savefig([pathst 'LogLogCurves_' pathst(end-15:end-1) '.fig']);

figure(1); legend(LegendStrings); savefig([pathst 'AllCurves_' pathst(end-15:end-1) '.fig']);

figure(3); yyaxis left; plot(times, slopes,'o');
yyaxis right; plot(times, shifts, 'o');
title(pathst);
savefig([pathst 'Slopes_' pathst(end-15:end-1) '.fig']);
end

function dist=CalcR2(shift, xvec, yvec, ind) % calculates the rsquared value
xvect=xvec-shift;
linfunc = @(p,x) p(1)*x-p(2); para0=[1 0];
[para1,~,resid,~,~,~,jacob] = lsqcurvefit(linfunc,para0,log(-xvect(yvec>10 & xvec<0)),log(yvec(yvec>10 & xvec<0)));
dist=(sum((linfunc(para1, log(-xvect(yvec>10 & xvec<0)))-log(yvec(yvec>10 & xvec<0))).^2)./sum((log(yvec(yvec>10 & xvec<0))-mean(log(yvec(yvec>10 & xvec<0)))).^2));
%(linfunc(para1, log(-xvect(yvec>10)))-log(yvec(yvec>10)));
end

%%measurement parameters for other experiments used in the study:

%%measurement of wing disc 2 with collagenase
% times=[-5, 0, 2, 7, 12, 17,22];
% pathst='~/mypath/ForceCurves_Drugs/WD_collagenase_20012021_2/'; % usable
% paths={[pathst '-5min_processed-2021.08.16-14.42.06/']; [pathst '0min_processed-2021.08.16-14.43.16/'];...
%     [pathst '0nismin_processed-2022.04.11-15.13.51/'];...
%     [pathst '5min_processed-2021.08.16-14.44.00/']; [pathst '10min_processed-2021.08.16-14.45.37/'];...
%     [pathst '15min_processed-2021.08.16-14.47.20/']; [pathst '20min_processed-2021.08.16-14.49.08/'];}
% Fmax=350; xmin=-1200; xmax=500;


%%measurement of wing disc 3 with collagenase
% times=[-5, 0, 2, 7, 12, 17];
% pathst='~/mypath/ForceCurves_Drugs/WD_collagenase_22012021/'; % usable
% paths={[pathst '-5min_processed-2021.08.16-14.21.17/']; [pathst '0min_processed-2021.08.16-14.22.52/'];...
%     [pathst '2min_processed-2021.08.16-14.25.53/']; [pathst '7min_processed-2021.08.16-14.28.21/'];...
%     [pathst '12min_processed-2021.08.16-14.23.46/']; [pathst '17min_processed-2021.08.16-14.30.32/']};
% Fmax=350; xmin=-800; xmax=500;
%

%%measurement of wing disc 4 with collagenase
% times=[0, 2, 7, 12, 17];
% pathst='~/mypath/ForceCurves_Drugs/WD72hAEL_collagenase_29012021/'; %usable, slopes increase
% paths={[pathst '0min_processed-2021.08.16-18.09.20/']; [pathst '2min_processed-2021.08.16-18.11.40/'];...
%     [pathst '7min_processed-2021.08.16-18.10.36/']; [pathst '12min_processed-2021.08.16-18.07.37/'];
% [pathst '17min_processed-2021.08.16-18.12.37/'];};
% Fmax=350; xmin=-800; xmax=500;

%%measurement of wing disc 5 with collagenase
% times=[-5, 17];
% pathst='~/mypath/ForceCurves_Drugs/WD_collagenase_22012021_2/'; %usable, slopes increase
% paths={[pathst '-5min_processed-2021.08.17-18.34.49/']; [pathst '17min_processed-2021.08.17-18.37.41/'];...
%         };
% Fmax=350; xmin=-1200; xmax=500;

%%measurement of wing disc 1 with latrunculin
% times=[-5, 17];
% pathst='~/mypath/ForceCurves_Drugs/WD_LatA_10032021_72hAEL/'; %usable, slopes increase
% paths={[pathst '72hAEL_1_0minprocessed-2021.08.18-10.11.46/']; [pathst '72hAEL_1_17minprocessed-2021.08.18-10.13.02/'];};
% Fmax=350; xmin=-800; xmax=500;

%%measurement of wing disc 2 with latrunculin
% times=[-5, 17];
% pathst='~/mypath/ForceCurves_Drugs/WD_LatA_10032021_2_72hAEL/'; %usable, slopes increase
% paths={[pathst '72hAEL_2_0minprocessed-2021.08.18-10.15.05/']; [pathst '72hAEL_2_17minprocessed-2021.08.18-10.19.30/'];};
% Fmax=350; xmin=-800; xmax=500;

%%measurement of wing disc 3 with latrunculin
% times=[-5, 17];
% pathst='~/mypath/ForceCurves_Drugs/WD_LatA_10032021_4_72hAEL/'; %usable, slopes increase
% paths={[pathst '72hAEL_4_0minprocessed-2021.08.18-10.20.57/']; [pathst '72hAEL_4_17minprocessed-2021.08.18-10.21.51/'];};
% Fmax=350; xmin=-800; xmax=500;
%
%%measurement of wing disc 4 with latrunculin
% times=[-5, 0, 2, 7, 12, 17];
% pathst='~/mypath/ForceCurves_Drugs/WD_LatA_02032021_72hAEL_2/';%conclusive
% paths={[pathst '72hAEL_2_-5minprocessed-2021.08.16-15.49.31/']; [pathst '72hAEL_2_0minprocessed-2021.08.16-15.50.34/'];...
%     [pathst '72hAEL_2_0bisminprocessed-2021.08.16-15.51.17/'];
%     [pathst '72hAEL_2_7minprocessed-2021.08.16-15.53.01/']; [pathst '72hAEL_2_12minprocessed-2021.08.16-15.53.41/'];...
%     [pathst '72hAEL_2_17minprocessed-2021.08.16-15.54.16/']};
% Fmax=350; xmin=-800; xmax=500;


%%measurement of wing disc 5 with latrunculin
% pathst='~/mypath/ForceCurves_Drugs/WD_LatA_05032021_72hAEL/';
% times=[-5, 0, 2, 7, 12, 17];
% paths={[pathst '72hAEL_1_-5minprocessed-2021.08.17-11.51.48/']; [pathst '72hAEL_1_0minprocessed-2021.08.17-11.59.03/'];...
%     [pathst '72hAEL_1_2minprocessed-2021.08.17-12.00.26/'];
%     [pathst '72hAEL_1_7minprocessed-2021.08.17-12.01.50/']; [pathst '72hAEL_1_12minprocessed-2021.08.17-12.03.16/'];...
%     [pathst '72hAEL_1_17minprocessed-2021.08.17-12.06.19/']; };
% Fmax=350; xmin=-800; xmax=500;
%
%%measurement of wing disc 6 with latrunculin
% pathst='~/mypath/ForceCurves_Drugs/WD_LatA_04032021_72hAEL/';%conclusive
% times=[-5, 0, 2, 7, 12, 17];
% paths={[pathst '72hAEL_1_-5minprocessed-2021.08.17-11.44.23/']; [pathst '72hAEL_1_0minprocessed-2021.08.17-11.45.17/'];...
%     [pathst '72hAEL_1_2minprocessed-2021.08.17-11.46.15/'];
%     [pathst '72hAEL_1_7minprocessed-2021.08.17-11.47.09/']; [pathst '72hAEL_1_12minprocessed-2021.08.17-11.48.12/'];...
%     [pathst '72hAEL_1_17minprocessed-2021.08.17-11.49.07/']};
% Fmax=350; xmin=-800; xmax=500;
%
%%measurement of wing disc 1 with Y27632
% pathst='~/mypath/ForceCurves_Drugs/WD_ROCK_11032021_72hAEL/';
% times=[-5, 0, 2, 7, 12, 17];
% % conclusive!!
% paths={[pathst '72hAEL_1_-5minprocessed-2021.08.16-18.38.25/']; [pathst '72hAEL_1_0minprocessed-2021.08.16-18.39.02/'];...
%     [pathst '72hAEL_1_2minprocessed-2021.08.16-18.42.46/'];
%     [pathst '72hAEL_1_7minprocessed-2021.08.16-18.41.47/']; [pathst '72hAEL_1_12minprocessed-2021.08.16-18.40.17/'];...
%     [pathst '72hAEL_1_17minprocessed-2021.08.16-18.41.07/']};
% % get 27 and 37 min
% Fmax=350; xmin=-800; xmax=500;
%
%%measurement of wing disc 2 with Y27632
% pathst='~/mypath/ForceCurves_Drugs/WD_ROCK_16032021_72hAEL/';
% times=[-5, 0, 2, 7, 12, 17,22];
% paths={[pathst '72hAEL_1_-5minprocessed-2021.08.17-11.04.24/']; [pathst '72hAEL_1_0minprocessed-2021.08.17-11.16.55/'];...
%     [pathst '72hAEL_1_2minprocessed-2021.08.17-11.18.16/'];
%     [pathst '72hAEL_1_7minprocessed-2021.08.17-11.20.40/']; [pathst '72hAEL_1_12minprocessed-2021.08.17-11.22.14/'];...
%     [pathst '72hAEL_1_17minprocessed-2021.08.17-11.23.29/'];[pathst '72hAEL_1_22minprocessed-2021.08.17-11.24.44/'];};
% %not much change at time points 37 and 47, took them out
% Fmax=350; xmin=-350; xmax=600;
%
%%measurement of wing disc 3 with Y27632
% pathst='~/mypath/ForceCurves_Drugs/WD_ROCK_26022021_72hAEL/';
% times=[-5, 0, 2, 7, 12, 17,27];
% paths={[pathst '72hAEL_2_BEFOREprocessed-2021.08.17-11.30.44/']; [pathst '72hAEL_2_BEFORE_0minprocessed-2021.08.17-11.31.53/'];...
%     [pathst '72hAEL_2_2minprocessed-2021.08.17-11.33.00/'];
%     [pathst '72hAEL_2_7minprocessed-2021.08.17-11.34.25/']; [pathst '72hAEL_2_12minprocessed-2021.08.17-11.35.22/'];...
%     [pathst '72hAEL_2_17minprocessed-2021.08.17-11.36.54/']; [pathst '72hAEL_2_27minprocessed-2021.08.17-11.37.52/'];};
% Fmax=350; xmin=-800; xmax=500;

%%measurement of wing disc 4 with Y27632
% pathst='~/mypath/ForceCurves_Drugs/WD_ROCK_17032021_72hAEL/';
% times=[ 0,  17];
% paths={[pathst '72hAEL_1_00minprocessed-2021.08.18-13.54.17/']; [pathst '72hAEL_1_17minprocessed-2021.08.18-13.55.01/'];};
% Fmax=350; xmin=-800; xmax=500;
```
